# Why Are Alzheimer’s Disease Patients Unaware of their Memory Deficits?

**DOI:** 10.1101/2022.02.01.478754

**Authors:** Solofo Razafimahatratra, Thomas Guieysse, François-Xavier Lejeune, Marion Houot, Takfarinas Medani, Gérard Dreyfus, André Klarsfeld, Nicolas Villain, Filipa Raposo Pereira, Valentina La Corte, Nathalie George, Dimitrios Pantazis, Katia Andrade

**Affiliations:** Institute of Memory and Alzheimer’s Disease, Salpêtrière Hospital, Paris, France; Sorbonne Université, Paris Brain Institute (ICM Institut du Cerveau), AP-HP, INSERM, CNRS, University Hospital Pitié-Salpêtrière, Paris, France.France; Paris Brain Institute’s Data and Analysis Core, University Hospital Pitié-Salpêtrière, Paris, France; Signal & Image Processing Institute, University of Southern California, Los Angeles, CA 90089, USA; ESPCI Paris – PSL, Paris, France; Laboratory of Brain Plasticity, CNRS UMR 8249, ESPCI Paris - PSL, Paris, France; Brain & Spine Institute, ICM, INSERM U 1127, CNRS UMR 7225, Sorbonne Université, Centre MEG-EEG, F-75013, Paris, France; McGovern Institute for Brain Research, Massachusetts Institute of Technology, Cambridge, MA, USA

## Abstract

Anosognosia, or the lack of awareness of one’s own impairment, is frequent for memory deficits in patients with Alzheimer’s disease (AD). Although often related to frontal dysfunctions, the neural mechanisms of anosognosia remain largely unknown. We hypothesized that anosognosia in AD may result from a failure in the error-monitoring system, thus preventing patients from being aware of and learning from their own errors. We therefore investigated the event-related potentials evoked by erroneous responses during a memory task in two groups of amyloid positive individuals who had only subjective memory complaints at study entry: 1) those who progressed to AD; and 2) those who remained cognitively normal after five years of follow-up. Our findings revealed direct evidence of a failure in the error-monitoring system at early stages of AD, suggesting that it may be the critical neural substrate of anosognosia in this neurodegenerative disorder.

## INTRODUCTION

Anosognosia for memory deficits is frequent in patients suffering from Alzheimer’s disease (AD), even at its early stage^1,2^. Typically, these patients are unaware of their memory impairment, which may have harmful consequences on their quality of life because it delays diagnosis and therapeutic interventions^2,3^, further increasing the burden of care^4^. An association of anosognosia with frontal dysfunctions has been found in AD patients, as revealed, for example, by executive deficits on the Trail Making Test^5^ and the Word Card Sorting Test^6^. In line with this, anosognosia in AD patients has been related to higher levels of apathy^3,7^. Nonetheless, to date, there is no gold standard for objective assessment of anosognosia, which instead relies on subjective measures of clinical judgment and discrepancy scores^8^. These typically include patient-caregiver rating discrepancies in distinct cognitive and functional domains, and the patient’s self-rating of her/his level of performance on a given task. Due to this heterogeneity of assessment methods, data about the prevalence of anosognosia in AD may vary considerably across studies^2^.

Neurocognitive models of anosognosia in AD have suggested either the involvement of motivational and executive control^9^ or comparator^10,11^ mechanisms. However, the precise characterization of the neural substrate of anosognosia is still unknown. Here, we hypothesized that anosognosia in AD may result from an inability to monitor ongoing behavior due to a failure in the error-monitoring system, which would normally signal the need for behavior adjustments to obtain an optimal performance^12,13^. Such impairment could also have adverse effects on global cognition by preventing subjects from learning from their own errors^14,15^.

Two event-related potentials (ERPs), namely the error-related negativity (ERN) and error positivity (Pe), have been established as the neural signatures of the executive system involved in error-monitoring^16,17^. The ERN is a negative deflection that peaks approximately 50 – 150 ms after the commission of an error (e.g., after an “incorrect” response in tasks that require the “correct” recognition of a stimulus). The Pe is a positive deflection following the ERN in a time window between 150 and 550 ms after an erroneous response. The earlier process of error detection is presumably implicit, whereas the later process of error awareness is considered explicit^18^. Accordingly, ERP results indicate that the Pe is associated with error awareness^19,20,21^, while the ERN is likely to reflect the activation of a more generic, preconscious system of error detection^16,17^ or conflict monitoring^22,23,24^, with additional evidence showing that the ERN is not always necessary for the emergence of the Pe and error awareness^25^. Some authors have further divided the Pe into an early (appearing until 400 ms after the ERN) and late subcomponents, suggesting that only the latter is modulated by error awareness^26^.

Source localization research indicates that both ERN and Pe are generated by the anterior cingulate cortex (ACC), with contributions from a more caudal region in the case of ERN^20,27^ and a more rostral region of the ACC, as well as the posterior cingulate cortex (PCC) in the case of Pe^20,28^. A prominent theory holds that ACC activity in response to errors might reflect a reinforcement learning process whereby subjects learn to associate actions with negative outcomes in order to improve goal-oriented behavior on the next occurrence^14^. Interestingly, the cingulate cortex is affected by amyloid deposition early on in aging and in AD^29^. However, although brain amyloid deposition confers a high risk for AD, the percentage of individuals with positive biomarkers who will progress to symptomatic clinical conditions is not yet known^30^.

Besides error monitoring, it has been suggested that these two ERP components may reflect other cognitive, emotional, and motivational processes^28^, which is consistent with evidence of equivalent evoked potentials occurring after correct responses^31,32^. These ERPs, called correct-related negativity (CRN) and correct positivity (Pc), have a similar topography and presumably the same brain origin as those evoked by errors^33^, suggesting that both CRN and Pc could be associated with brain processes identical to those implicated in the emergence of ERN and Pe, respectively, regardless of the related outcomes.

The main goal of this study was to assess the efficiency of the error-monitoring system in individuals at risk of developing AD. Specifically, we investigated whether a failure in the error-monitoring system of these individuals, as reflected by the ERN and the Pe amplitudes during a task of memory recognition of words, could distinguish those who progressed to AD at some point over the five-year study period, from those who remained cognitively normal all along. The study was based on a longitudinal cohort of subjects, who were cognitively normal at study entry despite subjective memory complaints^34^.

## RESULTS

A total of 51 subjects from the INSIGHT cohort^34^ who satisfied our eligibility criteria were included in this study. All individuals had an amyloid-positive status at study entry^35^. Out of the 51 individuals, 15 progressed to AD within the five years of the study duration (PROG group), and 36 remained cognitively normal (CTRL group). To assess the efficiency of the error-monitoring system, EEG data were recorded annually over five years while participants performed a memory task requiring the recognition of words seen between one and four hours before the EEG session. A failure to recognize whether a given word had been presented previously by the neuropsychologist, or not, constituted an error.

Demographic and neuropsychological characteristics of the PROG and CTRL groups are shown in Table 1. At entry (month M0), there were only minor, if any, cognitive differences between the groups in the score of the Mini Mental State Examination (MMSE)^36^ and the total recall score of the Free and Cued Selective Reminding Test (FCSRT)^37^. By contrast, differences on those neuropsychological tests became highly significant between the groups at Mdiag/M60 (i.e., either at month of AD diagnosis or at 60 months after entry). This reflected a decline of the PROG group and a relatively constant cognitive performance of the CTRL group over time. The demographic characteristics (age, gender and education level) did not differ significantly between groups. The Mdiag times were as follows: one subject converted to AD at M18, three subjects at M24, one subject at M36, one subject at M42, one subject at M54, and eight subjects at M60.

**Table 1.** Demographic and neuropsychological characteristics of the PROG and CTRL groups at M0 and Mdiag/M60. P-values were computed using a Wilcoxon-Mann-Whitney test for numerical variables (Age, MMSE) and Fisher’s exact test for categorical variables (Gender, Education level). Significant differences are indicated by asterisks: \*\*\**p* < 0.001, **0.001 ≤ *p* < 0.01, *0.01 ≤ *p* < 0.05. ^a^ Categorization of the education level into “high” and “low or intermediate” education levels are as indicated in Dubois et al., 2018.^34^

| Population<br>Characteristics | M0 |  |  | Mdiag/M60 |  |  |
| --- | --- | --- | --- | --- | --- | --- |
|  | PROG<br>n=15 | CTRL<br>n=36 | p-value | PROG<br>n=15 | CTRL<br>n=36 | p-value |
| Gender (M/F, % F) | 7/8, 53% | 14/22, 61% | 0.76 |  |  |  |
| Age<br>mean ± SD (range) | 77.5 ± 3.4<br>(70 – 85) | 76.3 ± 3.0<br>(71 – 84) | 0.18 | 81.7 ± 3.2<br>(75 – 86) | 81.3 ± 3.0<br>(76 – 89) | 0.44 |
| Education level <sup>a</sup><br>High/Low-Intermediate,<br>% High | 10/5, 67% | 17/19, 47% | 0.23 |  |  |  |
| MMSE (maximum 30)<br>mean ± SD (range) | 28.2 ± 0.7<br>(27 – 29) | 28.7 ± 0.9<br>(27 – 30) | 0.07 | 25.9 ± 3.9<br>(14 – 30) | 28.5 ± 1.5<br>(23 – 30) | 0.002** |
| FCSRT-Total Recall<br>(maximum 48)<br>mean ± SD (range) | 44.7 ± 2.2<br>(41 – 48) | 45.9 ± 1.8<br>(41 – 48) | 0.043* | 32.5 ± 8.7<br>(10 – 41) | 45.5 ± 2.2<br>(40 – 48) | <<br>0.001*** |

### Behavioral performance on the memory task

We compared the overall performance of the participants on the memory recognition of words task in terms of accuracy (percentage of correct responses over 208 trials) and reaction time (RT). To this end, two linear mixed models (LMMs) were fitted to investigate the effects of group (PROG; CTRL), time (M0; Mdiag/M60), type of response (Correct or Incorrect/Error, only for RT), and all interaction terms. Both models included a random intercept on the subject identifier and two covariates for age and education level at M0. Main and interaction effects (“omnibus” test) of the fitted LMMs are reported with Wald χ^2^ statistic and degrees of freedom (df); and post hoc pairwise comparisons (including a Benjamini-Hochberg correction of the false discovery rate) on significant main and interaction effects are reported with t-ratio statistic (t) and Kenward-Roger’s approximation for the degrees of freedom. More information on the statistical analyses is provided in the Methods section.

Both LMMs robustly distinguished the PROG and CTRL groups in terms of accuracy and RT (Figures 1 and 2, respectively; also see Supplementary Tables 1 and 2 for detailed list of effects). Critically, Type II Wald chi-square tests for the two fitted models indicated a significant “Group × Time” interaction effect on accuracy (*p* <0.001) and RT (*p* <0.001). A further investigation of this interaction with pairwise comparisons based on estimated marginal means ± standard error (SE) revealed that the PROG group was significantly less accurate (*p* <0.001) at Mdiag/M60 (78.9 ± 2.7 %) than at M0 (93.4 ± 1.6 %), indicating a decrease of accuracy over time. By contrast, the accuracy of the CTRL group remained high between these two time points. As a result, the two groups ended up differing significantly (*p* <0.001) at Mdiag/M60, where the accuracy was worse for the PROG group (78.9 ± 2.7 %) than for the CTRL group (97.5 ± 0.7 %).

**Figure 1.**
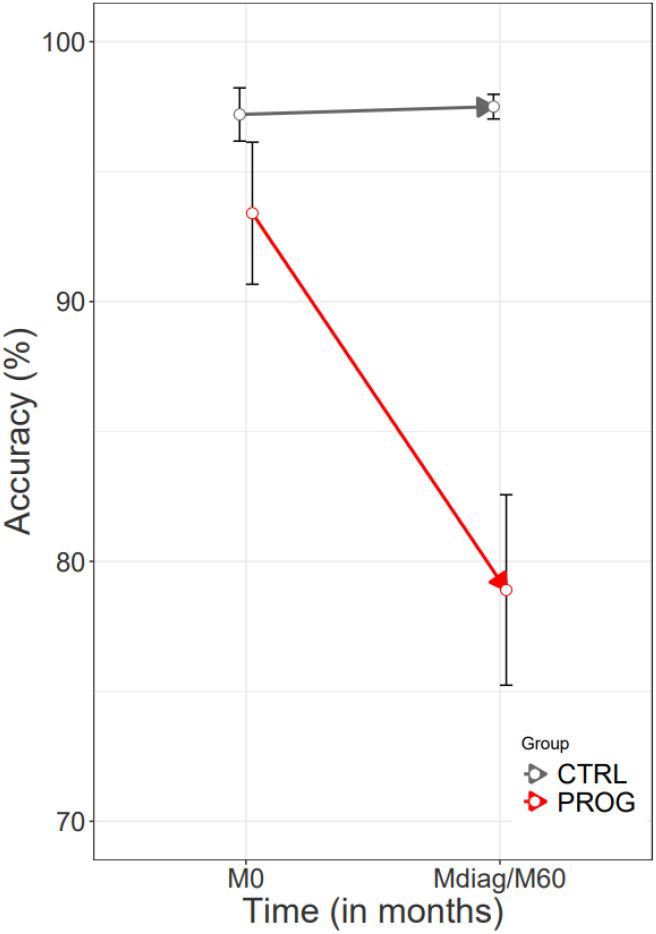
Behavioral accuracy of the PROG and CTRL groups on the memory task at times M0 and Mdiag/M60. Plots show percentage of correct responses together with 95% confidence intervals.

**Figure 2.**
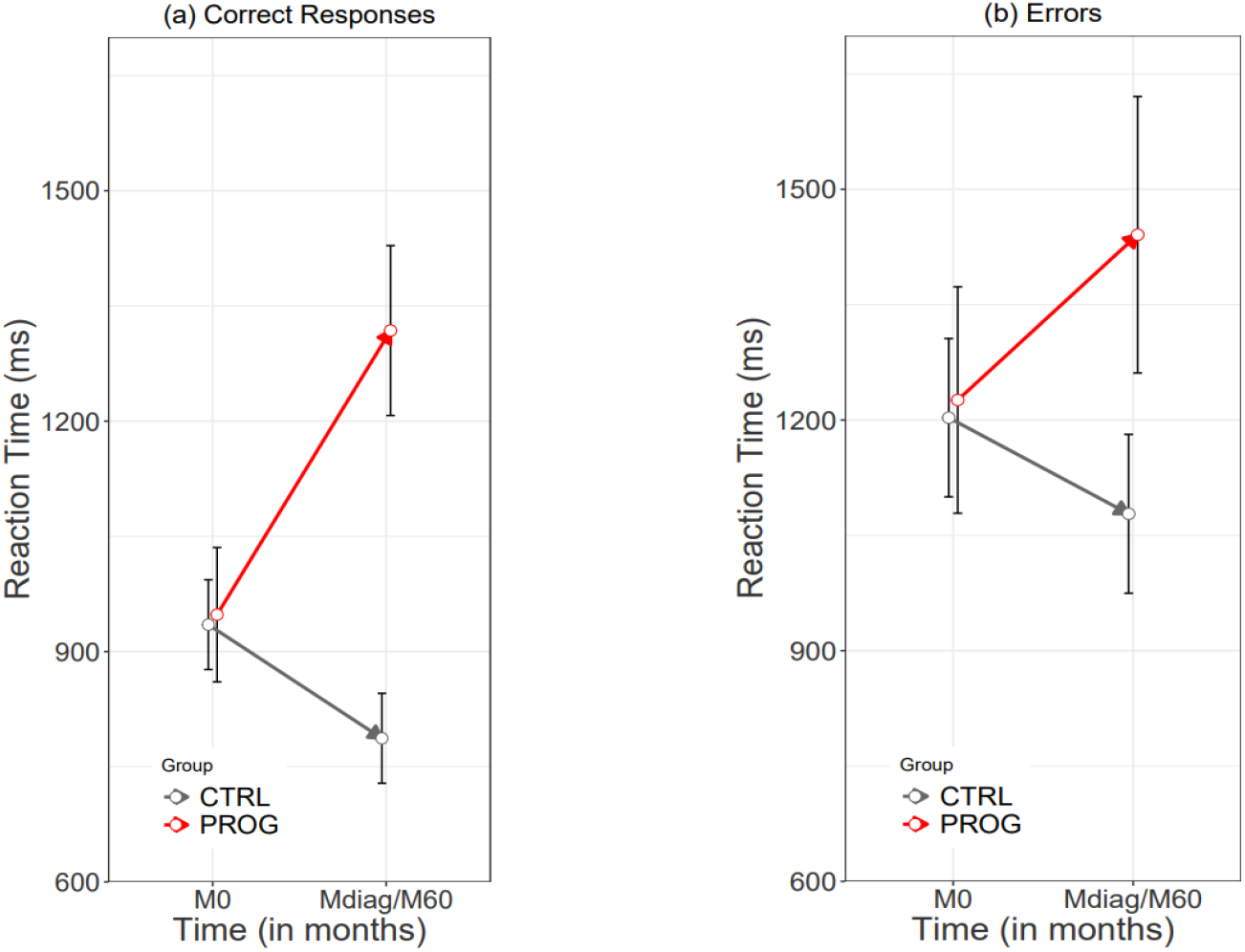
Reaction times of the PROG and CTRL groups on the memory task at times M0 and Mdiag/M60. Plots show the mean of reaction times with 95% confidence intervals for (a) correct responses, and (b) errors.

The RT results were consistent with these findings. Specifically, the PROG group had significantly longer RT (*p* <0.001) at Mdiag/M60 for all responses (correct and erroneous combined, 1380 ± 112 ms) than at M0 (1087 ± 113 ms), which further resulted in the two groups differing significantly (*p* <0.01) at Mdiag/M60 regardless of the type of response. This difference became markedly larger due to the fact that the RT of the subjects of the CTRL group decreased significantly (*p* < 0.05) for all responses (correct and erroneous combined, 1069 ± 72 ms at M0 versus 933 ± 72 ms at Mdiag/M60) between the two time points (as detailed in Supplementary Table 1b). Furthermore, we observed that the response times for erroneous responses (1237 ± 66 ms) were significantly larger than those for correct responses (998 ± 65 ms) over both groups and both time points (*p* < 0.001) (Figure 2).

Taken together, these results revealed a learning disability emerging in the PROG group from M0 to Mdiag, which clearly contrasted with the subjects of the CTRL group, whose behavioral performance remained relatively unchanged (accuracy) or even improved (reaction time) over time.

### ERP analysis on errors

For reliable ERP estimation, we required a minimum of six artifact-free trials for each subject and response type (Correct or Incorrect/Error)^38^. This led to reduced sample sizes of n=9 at M0 and n=15 (unchanged) at Mdiag for the PROG group, and n=10 at M0 and n=9 at M60 for the CTRL group for the EEG analyses. Demographic and neuropsychological characteristics of these subsamples of subjects are shown in Supplementary Table 3.

For ERP analysis, we fitted four LMMs to separately model the two ERP components (ERN and Pe, where the latter was divided into two early and two late subcomponents) at each electrode location (FCz and Cz). The LMMs assessed the effects of group (PROG, CTRL), time (M0, Mdiag/M60), two early and two late subcomponents for Pe, and all the resulting interaction terms. All models included a random intercept on the subject identifier and two covariates for age and education level at M0. The ERN component was estimated as the mean amplitude within −15/+15 ms around the most negative peak in the time interval from 0 to 150 ms after an erroneous response^39^. The Pe for each subcomponent was estimated as the mean amplitude for 100 ms time windows from 150 to 550 ms^26^.

The ERN and Pe responses for the PROG and CTRL groups are shown on Figure 3. The LMM results revealed a significant interaction effect of “Group × Time” on Pe at both FCz (*p* < 0.001) and Cz (*p* < 0.001) electrodes, thus indicating that group differences are time-dependent. There were no significant differences between early and late Pe subcomponents, thus suggesting that both may reflect the same neural mechanism. A detailed description of the LMM results is presented in Supplementary Tables 4 and 5.

**Figure 3.**
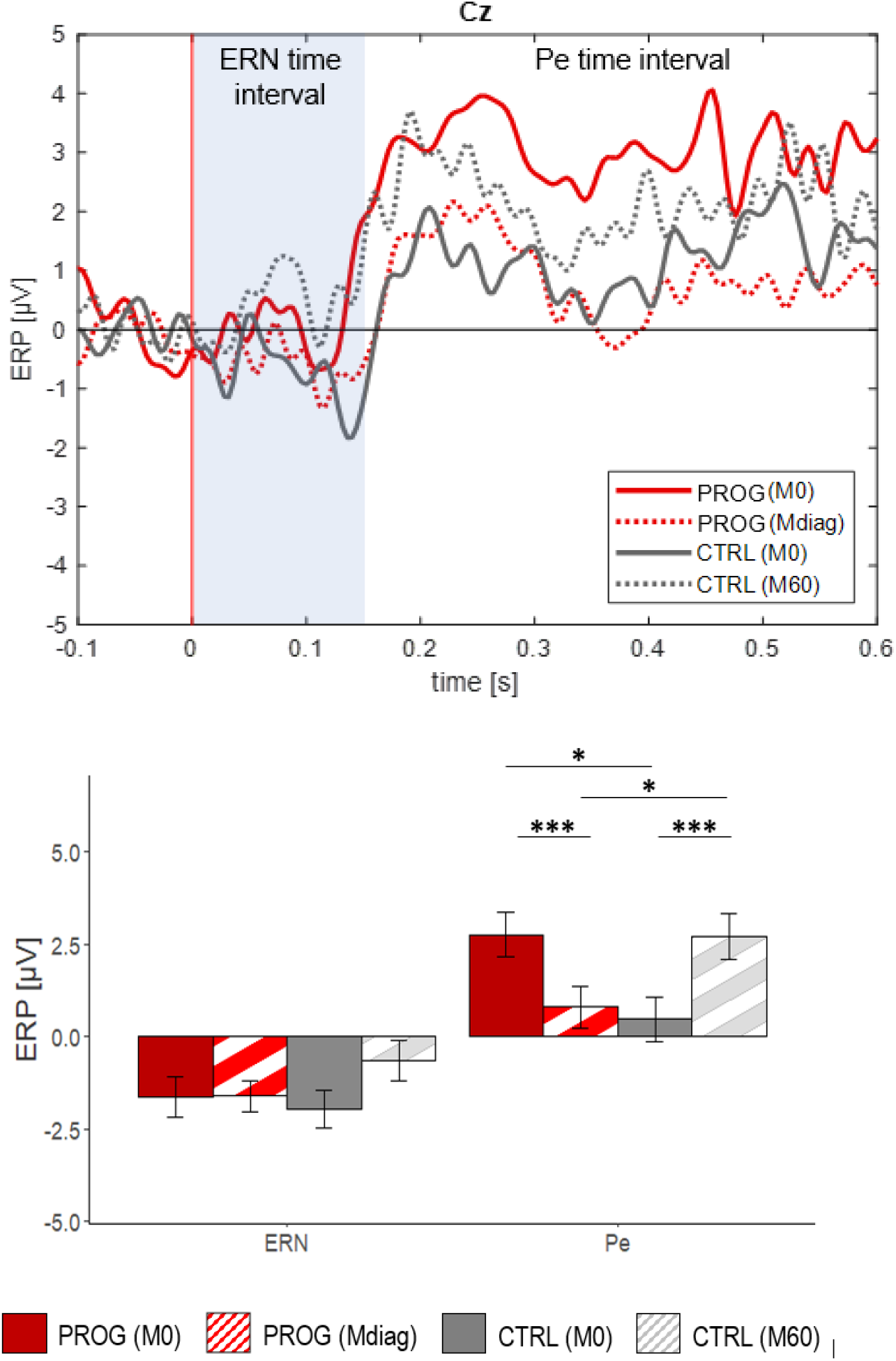
Grand average ERP (ERN and Pe) amplitudes, time-locked to the (erroneous) response occurring at time 0 s, recorded at electrode Cz. ERN is found in the 0 – 150 ms time interval post-error (top panel). Pe is found within the 150 – 550 ms time interval. Barplots show the estimated marginal means and standard errors of the ERPs at M0 and Mdiag/M60 for both groups (bottom panel). Significant differences are indicated by asterisks: \*\*\**p* < 0.001, **0.001 ≤ *p* < 0.01, *0.01 ≤ *p* < 0.05

From the “Group × Time” interaction, the following post hoc comparisons on estimated marginal mean amplitudes (± SE) showed direct evidence of a failure in the error-monitoring system during a memory task in early stages of AD. Specifically, at Cz electrode, the Pe amplitude decreased significantly from M0 to Mdiag for the PROG group (2.75 ± 0.61 μV at M0 versus 0.79 ± 0.57 μV at Mdiag, *p* < 0.001), whereas the CTRL group showed the inverse pattern of evolution (0.47 ± 0.61 μV at M0 versus 2.71 ± 0.63 μV at M60, *p* < 0.001), as illustrated in Figure 3 A similar evolution was found at FCz electrode, although with lower Pe amplitudes for both groups: (0.61 ± 0.49 μV at M0 versus 0.14 ± 0.45 μV at Mdiag, *p* = 0.145, for the PROG group; and CTRL −0.31 ± 0.48 μV at M0 versus 1.84 ± 0.50 μV at M60, *p* < 0.001, for the CTRL group) (Figure 4). Importantly, these results strongly suggest that different phenomena reflected in the Pe amplitude^40^ may contribute to distinguishing the two groups between M0 and Mdiag/M60: 1) a lack of awareness for memory impairment (erroneous responses in the memory task) at Mdiag/M60 in those subjects who progressed to AD (i.e., the PROG group); and 2) a learning effect, probably resulting from an improved ability to recognize erroneous responses and to adjust one’s behavior with the repetition of the task, in those subjects who remained cognitively normal over time (i.e., the CTRL group).

**Figure 4.**
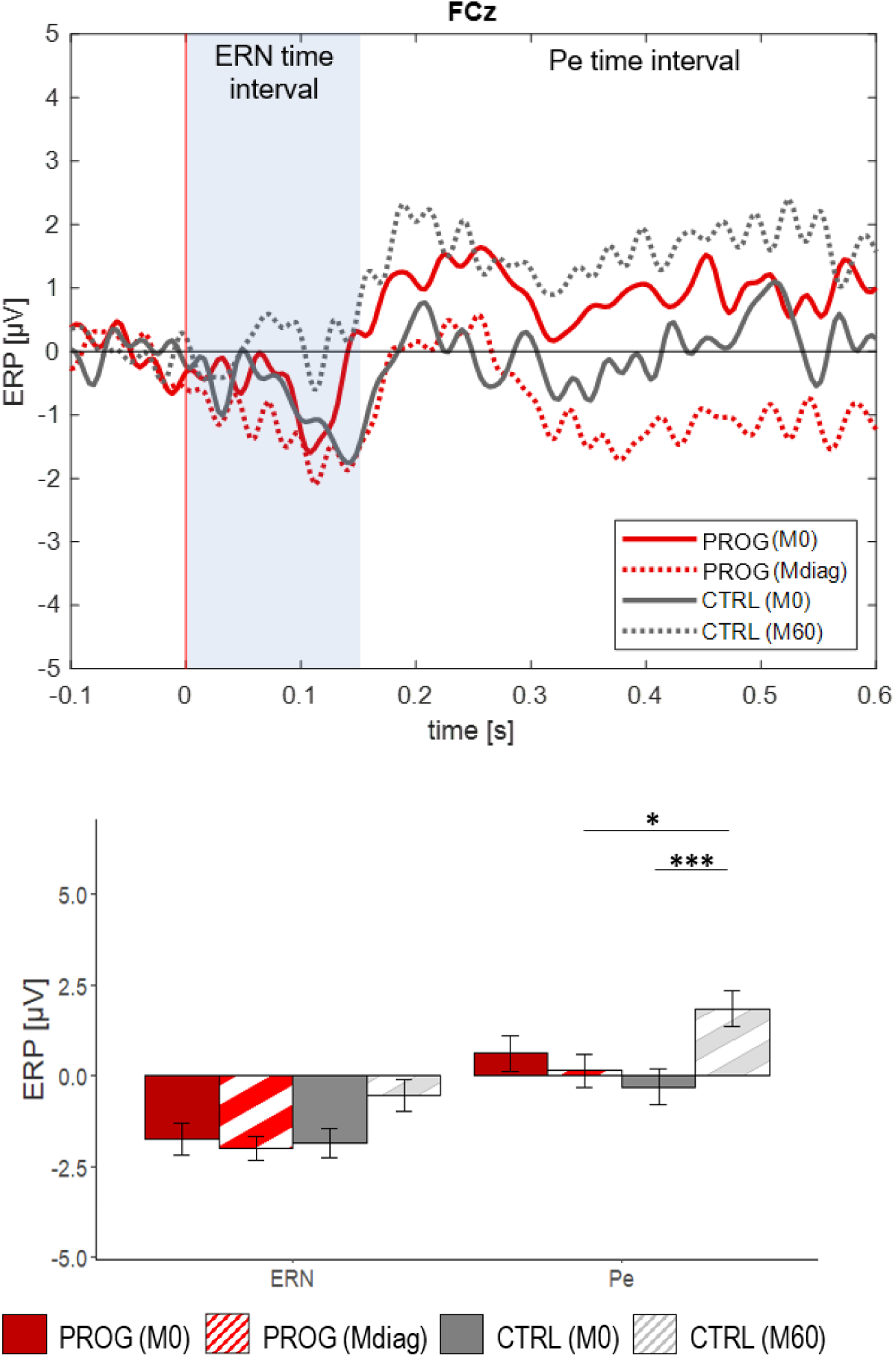
Grand average ERP (ERN and Pe) amplitudes, time-locked to the (erroneous) response occurring at time 0 s, recorded at electrode FCz. ERN is found in the 0 – 150 ms time interval post-error (top panel). Pe is found within the 150 – 550 ms time interval post-response. Barplots showing the estimated marginal means and standard errors of ERPs at M0 and Mdiag/M60 for both groups (bottom panel). Significant differences are indicated by asterisks: \*\*\**p* < 0.001, **0.001 ≤ *p* < 0.01, *0.01 ≤ *p* < 0.05

Interestingly, at M0, the Pe amplitude of the PROG group was significantly higher than the Pe amplitude of the CTRL group at Cz electrode (0.47 ± 0.61 μV at M0 for the CTRL group versus 2.75 ± 0.61 μV at M0 for the PROG group, *p* < 0.05), further suggesting the existence of a neural compensation mechanism that probably helped the subjects of the PROG group to accurately perform the memory task at entry (that is, up to five years before their AD diagnosis).

At FCz, where the ERN is typically maximal, we observed that the magnitude (absolute value of the amplitude) of the ERN for the CTRL group at M60 was lower than for the PROG group at Mdiag (0.54 ± 0.44 μV at M60 versus 2.00 ± 0.34 μV at Mdiag, *p* = 0.058), but it remained only a trend. An additional trend was found for a decrease of the ERN magnitude between M0 and M60 in the CTRL group (1.85 ± 0.42 μV at M0 versus 0.54 ± 0.44 μV at M60, *p* = 0.073), while no differences on the ERN were found between these two time points in the PROG group (1.76 ± 0.44 μV at M0 versus 2.00 ± 0.34 μV at Mdiag, *p* = 0.87). At Cz electrode, neither significant effect nor even a trend was found on the ERN (thus no post hoc comparison was performed). Details of the post hoc tests are reported in Supplementary Table 3.

### ERP analysis on correct responses

For ERP analysis on correct responses, four separate models were constructed in a similar manner to those based on the erroneous response data for the two respective ERP components (CRN and Pc) within the same time intervals as above, at the FCz and Cz electrodes.

As detailed in Supplementary Table 5, Type II Wald chi-square tests for the fitted model indicated a significant effect of “Group × Time” on Pc models at both Cz (*p* < 0.001) and FCz (*p* < 0.001) electrodes. No significant effect of the CRN component was found. Post hoc comparisons on the “Group × Time” interaction showed a significant decrease of the Pc amplitude from M0 to Mdiag for the PROG group at Cz (1.65 ± 0.40 μV at M0 versus 0.85 ± 0.37 μV at Mdiag, *p* < 0.01); and at FCz (0.62 ± 0.27 μV at M0 versus 0.19 ± 0.24 μV at Mdiag, *p* < 0.05). By contrast, the CTRL group showed a significant increase of the Pc amplitude from M0 to M60 at the same fronto-central sites (Cz, 0.72 ± 0.39 μV at M0 versus 2.30 ± 0.41 μV at M60, *p* < 0.001; and FCz, 0.19 ± 0.27 μV at M0 versus 1.31 ± 0.28 μV at M60, *p* < 0.001). Moreover, a significant difference was found between the two groups at Mdiag/M60 also at Cz (2.30 ± 0.41 μV at M60 versus 0.85 ± 0.37 μV at Mdiag, *p* < 0.05); and at FCz (1.31 ± 0.28 μV at M60 versus 0.19 ± 0.24 μV at Mdiag, *p* < 0.05). This pattern of evolution was similar to that of the Pe amplitude for the same subsamples of subjects as presented above in the respective subsection. Details of the post hoc tests are reported in Supplementary Table 4.

## DISCUSSION

Episodic memory relies on constructive processes that are sometimes prone to error^41^. Learning from our previous mistakes, avoiding the repetition of errors, requires an error-monitoring system that allows for rapid detection and evaluation of errors in order to adjust our behavior or improve our performance on goal-directed tasks^12^. This ability might be necessary for subjects to identify an error when unable to distinguish between “true” and “false” episodic memories. An amnestic episodic syndrome is typically observed in prodromal stages of AD^42^. The hippocampus is primarily involved in the formation of this type of memory, but also in episodic retrieval, a process in which other brain regions participate, including the prefrontal cortex and the PCC^43^.

At Mdiag/M60, the reaction times and Pe amplitudes of the PROG group were significantly smaller, and the error rates larger, than they were at M0, and than those of the CTRL group at M60. Consistent with the ERP results on errors, EEG potentials evoked by correct responses followed a similar pattern of evolution over time within and between groups. In association with the behavioral results on accuracy and reaction time, these ERP findings seem to reflect a lack of awareness for memory deficits, as well as a learning disability of the subjects of the PROG group at the moment of their AD diagnoses. Notably, the Pe, but not the ERN, seems a robust neural index of error awareness, with additional evidence of post-error behavioral modulation by consciously perceived errors^44^. In line with this, it has been proposed that an increased Pe could reflect a learning strategy allowing subjects to avoid future errors^39^, with further assumptions that both CRN and Pc could be associated with brain processes identical to those implicated in the emergence of ERN and Pe, respectively.

Taken together, our ERP results strongly suggest that an early impairment in the error-monitoring system of AD patients may critically contribute to their amnestic syndrome by preventing patients from consciously identifying “true” versus “false” memories (that is a clear sign of anosognosia) and learning from their own errors – inabilities that could explain why Alzheimer’s disease patients are unaware of their memory deficits. Supporting this view, recent neuroimaging studies have shown a reduced glucose metabolism in both the PCC^45,46^ – which is one of the brain sources of the Pe^20^ – and the hippocampus^46^ in prodromal AD patients with anosognosia for memory deficits. As a result, a decrease in Pe amplitude evoked by erroneous responses during an episodic memory task might constitute an objective biomarker of this kind of anosognosia in patients with AD and may therefore contribute to its identification from the very early stages of this neurodegenerative disorder.

On the other hand, the higher amplitude of the Pe at M0 than at Mdiag in the PROG group suggests a neural mechanism of “less-wiring-more-firing” type^47^, probably activated in these subjects in order to perform the memory task with success despite some difficulties observed already at study entry as reflected by a normal, although lower total recall score on the FCSRT when compared to the subjects of the CTRL group at the same time point (Supplementary Table 3). Accordingly, evidence from monkey studies has shown an increase in the PCC firing rate early in conditions of poor learning related, for example, to the lack of focus during an experiment^48^. Taking into account its origin^20^, it seems very plausible that the higher Pe amplitude observed at M0 in the PROG group reflects a similar neural mechanism, allowing these subjects to still compensate for subclinical difficulties in learning. Such a neural marker may help to distinguishing between subjects who will progress or not to AD in a period of five years and may therefore contribute to identify preclinical stages of AD and facilitate research trials for secondary prevention in this neurological domain^30^.

Lastly, whether the ERN and the Pe represent two independent error-monitoring processes is still an open question^25^. According to our results, the amplitude of the ERN does not necessarily co-vary with the amplitude of the Pe. In fact, while the Pe amplitude – particularly related to error awareness – decreased over time for the PROG group (and increased for the CTRL group), the ERN amplitude did not change significantly between the two studied time points for both groups.

The ERN has been associated with preconscious error-monitoring processes and may vary with stimulus novelty^49^ or expectedness^50^, with evidence of higher ERN magnitudes (i.e., more negative) occurring after unexpected stimuli than after expected ones^13^. Due to their memory deficits, it is possible that the “surprise effect” or novelty of the presented words (that is, the task stimuli) remained relatively intact for the PROG group at Mdiag and, therefore, without impact on the ERN amplitude. For the CTRL group, it is possible that the practice effect resulting from the task repetition over time had contributed to the reduction of the ERN magnitude at M60^50^, although this finding remained only a trend. Overall, the results on the ERN reinforce the hypothesis that the ERN and the Pe are at least partly independent and may indeed reflect distinct brain functions implicated in performance monitoring and self-awareness.

Despite the relatively reduced sample sizes for ERP analysis, our results on both correct and erroneous responses are consistent within and between groups at distinct time points (M0; Mdiag/M60) in two fronto-central sites (FCz; Cz).

## CONCLUSION

The present work is based on an original approach focused on the study of the error-monitoring system by combining behavior and ERP data analysis in a longitudinal cohort of individuals at risk for AD.

Our findings provide direct evidence of a failure in the error-monitoring system during a memory task from the early stages of AD, suggesting that it may be the critical neural substrate for the emergence of anosognosia in this neurodegenerative condition by preventing patients to be aware of their own memory impairment. This seems particularly corroborated by a significant decrease in the Pe amplitude in the subjects of the PROG group between M0 and Mdiag, which is consistent with a lack of awareness for memory deficits often observed in AD patients in the clinical setting.

Overall, in association with the neural compensation mechanism observed at entry (that is, up to five years before AD diagnosis), the results of this study allow us to propose that an early failure in the error-monitoring system may be not only the neural substrate of anosognosia in AD, but also a clue to future AD progression among individuals with a positive-amyloid status. This is important in light of an anticipatory medicine that would be of particular interest to deal with this devastating neurological disorder in the near future.

## METHODS

### Participants

We focused this study on two subgroups of the INSIGHT cohort^34^ identified after five years of follow-up, both of which presented at entry a brain pathological amyloid status evaluated by 18F-florbetapir (AV-45) positron emission tomography (PET) imaging: 1) amyloid-positive subjects who progressed to AD (PROG group); and 2) amyloid-positive subjects who did not progress to AD (CTRL group). For amyloid PET images, standard uptake value ratios were calculated by averaging the mean activity of cortical regions of interest: both left and right precuneus, cingulum posterior, cingulum anterior, and parietal, temporal and orbitofrontal cortex (the reference region was a combination of whole cerebellum and pons). The threshold set for normal (amyloid negative) versus abnormal (amyloid positive) uptake is 0.7918^35^. A second inclusion criterion was a FCSRT-total recall score < 47 at least once at M0 or M60 (our two main points of interest) to approximately balance the memory performance across subjects of the CTRL group.

A total of 88 out of 318 healthy volunteer subjects of the INSIGHT cohort were amyloid-positive at study entry (M0). Of these, 19 did not have behavioral/neural data at M60, and 18 had a high FCSRT^37^ total recall score (≥ 47) at both M0 and M60. Thus, our sample comprised 51 individuals, of which 15 progressed to AD within the 5 years of the study duration, and 36 remained cognitively normal. For behavioral data, the number of subjects was n=15 for the PROG group and n=36 for the CTRL group. For EEG data, we required a minimum of six artifact-free trials for both correct and erroneous responses (see EEG signal recording and pre-processing section below) at M0 and at Mdiag/M60. Therefore, given the relatively scarce error responses by some individuals, for EEG data analysis the number of subjects was n=9 at M0 and n=15 at Mdiag for the PROG group, and n=10 at M0 and n=9 at M60 for the CTRL group, as shown in Supplementary Table 3.

Descriptive characteristics for these two groups included demographic and clinical data on age, sex, education level (indicated by the scoring adopted by Dubois et al., 2018, covering pre-primary to tertiary levels of education on a scale ranging from 1 to 8; scores ≥ 7 correspond to “high” education levels and scores ≤ 6 indicate “lower or intermediate” education levels), the MMSE score^36^ and the total recall score of the FCSRT^37^ at M0 (i.e., at entry, month M0) and Mdiag/M60 (i.e., at month of AD diagnosis or 60 months from study inclusion, as previously mentioned). Importantly, there is evidence that the amnestic syndrome identified by the FCSRT can distinguish patients at an early stage of AD from mild cognitive impairment non-converters^51^, as well as AD from non-AD dementias^52^.

Data are presented in Table 1 and Supplementary Table 3 as mean ± standard deviation (SD) and range (minimum and maximum) for numerical variables, and frequency counts and percentages for categorical variables. Wilcoxon-Mann-Whitney test for numerical variables or Fisher’s exact test for categorical variables were used to compare the groups at each time point (M0 or Mdiag/M60).

### Experimental design

EEG data were collected annually for each participant over a five-year period while the participants performed a cognitive task of memory recognition of words: participants were presented with sequences of words, and instructed to indicate which of them they had previously memorized.

Precisely, the stimuli consisted of 16 target words sampled from the FCSRT^37^, and 64 distractor words that matched the target words in frequency of occurrence and length. The participants were asked to memorize the study list (target words) between one and four hours before the EEG session.

Each trial started with a fixation cross displayed in the center of the screen for 1.5 s, followed by the presentation of the stimulus (word item) for 2 s. Then a question prompted the participant to indicate whether the word was one of the target words (yes/no, corresponding to target and distractor words, respectively) by pressing buttons in a two-alternative forced choice design. No feedback was provided to the subjects on their performances. The first question was followed by a second one: “Are you sure of your answer?”. This question also required a binary response (yes/no), which indicated whether subjects were in doubt about their answers (certainty, yes; versus uncertainty, no), without quantifying their level of doubt (or uncertainty). The present study focused only on the subjects’ responses to the first question in terms of behavior and EEG potentials evoked by correct or erroneous answers during the described memory task. In particular, the experiment began with a training phase, during which participants performed a short training block (1min duration) to familiarize them with the EEG task setup. The episodic memory test then consisted of four successive blocks of 52 trials each, for a total of 208 trials. These trials presented in pseudorandom order each target word five times and the distractor words as follows: 16 distractors were presented five times, while the remaining 48 were presented only once during the all experiment (i.e., the four blocks). These blocks were separated by a break, with the task proceeding to the next block when the participant signaled she/he was ready. Stimulus presentation was controlled by E-prime software version 2.0 (Psychology Software Tools, Pittsburgh, PA), with words being displayed in white color on a black screen monitor at 1 meter distance from the participant. The episodic memory test lasted about 20 minutes.

A failure to recognize whether a given word had been presented previouslyby the neuropsychologist, or not, constituted an error (i.e., either a false recognition of a distractor word or an omission of the recognition of a target word). Individuals were not asked to respond quickly, which could have increased the number of errors (see Bogacz et al,^54^ for a review on the speed-accuracy tradeoff), but were allowed to respond as soon as they felt ready, with no upper time limit set for response. Behavioral data in this experiment consisted of the subjects’ accuracy (percentage of correct responses over 208 trials per subject) and the time interval between the presentation of the stimulus and their responses (reaction time; RT). Importantly, RT could decrease even after accuracy reached 100%, thus providing valuable behavioral assessment even in situations in which task performance is relatively error free. Both accuracy and RT were then investigated as behavioral clues to memory and learning abilities.

### EEG signal recording and pre-processing

EEG data were acquired with a 256-channel whole-head Geodesic 300 EEG System (Electrical Geodesics, Eugene, OR, USA) at a sampling rate of 250 Hz. Electrooculogram (EOG) data were recorded from electrodes placed above, below, and lateral to each eye. All electrode impedances were kept below 50 kΩ. The initial reference electrode was placed at Cz site. Sensor layout for 256-channel Hydrocel Geodesic Sensor Net is given in Supplementary information.

The Brainstorm software was used for pre-processing (Tadel et al., 2011). As the channel Cz was of interest for analyzing the ERN and the Pe amplitudes, we first re-referenced the EEG signals from Cz to the mean of the Left and Right Mastoids channels (CH094/CH190) (cf. Supplementary Figure 1). The data were then band-pass filtered between 0.1 and 40 Hz. Blinks were automatically detected from four pairs of channels (CH018-CH037, CH025-CH032, CH010-CH046, CH033-CH019; cf. Supplementary Figure 1). Four steps of blink detection were performed consecutively for each of the pairs previously cited. Blinks detected from a pair of channels are grouped in one series. Based on each blink series, the Signal-Space Projection (SSP) algorithm was applied separately to remove blink artifacts leading to four projectors. The projector on which the artifacts were most clearly detected was retained. After artifact suppression (from SSP), trials were extracted in the time interval of [-100 ms, 600 ms] with respect to the button press response time. Individual trials containing residual blink-related artifact when compared before and after component suppression or high amplitude variation (peak-to-peak > 100 μV) were detected visually and rejected..

For electrodes Cz (CH257) and FCz (CH015), which were our channels of interest, ERPs were estimated from the averaged trials after removing the baseline mean activity of each channel in the 100 ms window prior to response. Specifically, the ERN was determined as the mean amplitude within −15/+15 ms around the most negative peak in the time interval from 0 to 150 ms after an erroneous response^39^. The Pe was subdivided into two early (Pe1, 150–250 ms; Pe2, 250–350 ms) and two late (Pe3, 350–450 ms; Pe4, 450–550 ms) subcomponents, being determined for each subcomponent as the mean amplitude for 100ms time windows from 150 to 550 ms^26^. The same time intervals were used for their correct-response counterparts, CRN and Pc.

We required a minimum of six artifact-free trials on errors (or correct responses) for a reliable ERP estimation. This is in line with previous findings showing that six to eight error trials are sufficient to achieve adequate consistency for ERN and Pe amplitudes^38^. As a result, EEG data analysis was conducted on subsamples of subjects as mentioned in the “Participants” section (see above).

### Statistical analyses

All statistical analyses were conducted using R version 3.6.1 (R Development Core Team)^55^ and plots were generated with the ggplot2 package (v3.3.2)^56^. The level of statistical significance was set to *p* < 0.05 for all tests.

Within and between group differences and changes were examined using linear mixed-effects models (LMMs). Specifically, models were built for each outcome of interest in the behavioral (two models, one for accuracy and the other for RT) and the EEG (eight models, for the amplitudes of the ERN/CRN and Pe/Pc components at FCz and Cz electrodes, respectively) data. In the behavioral models, the factors Group (PROG, CTRL), Time (M0, Mdiag/M60), and Type of response (Correct, Error) only for RT, were regarded as fixed effects. We further included the interaction terms between these factors as fixed effects. In the ERP models related to either correct or erroneous responses, fixed effects involved Group (PROG, CTRL), Time (M0, Mdiag/M60), the addition of the two early and two late Pe/Pc subcomponents for the Pe/Pc models, and all the resulting interaction terms. The subject identifier was assigned as a random effect (intercept) to account for the repeated measurements. Age and education level at M0 were also included for covariate adjustments. All LMMs were fitted using restricted maximum-likelihood estimation (REML) from the function lmer in the lme4 package (v1.1-21)^57^. Significance for the main effects and all two-way and three-way interactions was assessed based on Type II Wald chi-square tests using the function Anova in the car package (v3.0-7). Post hoc pairwise comparisons were carried out on significant higher interaction with the emmeans package (v1.4.5) allowing the specification of custom contrasts between specific group means of interest to further determine where the differences occurred. P-values resulting from the post hoc tests were determined from the t-ratio (t) with Kenward-Roger’s approximation for degrees of freedom (df), and after adjustment by the Benjamini and Hochberg false discovery rate procedure to account for the multiplicity of contrasts. Results of the post hoc tests were also reported with the estimated marginal means (i.e., means predicted by the LMMs) and standard errors (SE) of the groups compared using emmeans. For the behavorial accuracy model, an arcsine square root transformation was applied to the proportion data prior to modeling as this improved the model assumptions of linearity, normality and constant variance of residuals. Finally, all significant interaction effects were also confirmed with Type III Wald chi-square tests using the function Anova in the car package (v3.0-7).

## ACKNOWLEDGMENTS

The authors wish to thank Prof. Bruno Dubois for his helpful comments and invaluable research support. Our acknowledgment extends to the INSIGHT-preAD cohort group (to which KA also belongs).

## AUTHOR CONTRIBUTIONS

KA conceived the hypothesis, designed and conducted the study; participated in the acquisition of clinical data; wrote the first draft and the final manuscript.

SR and TG performed behavioral and EEG analysis since the preprocessing step, performed the statistical analyses, discussed the results and contributed to the text, tables and artwork of the final manuscript.

FXL and MH conducted the statistical analysis and contributed to the writing of the final manuscript. FXL also contributed to the tables included in Supplementary information.

TM co-supervised EEG preprocessing and contributed to the discussion of the results and to the writing of the final manuscript.

GD and AK contributed to the analysis and discussion of the results and to the writing of the final manuscript. GD further contributed to the preliminary statistical analysis.

FRP, NV, VLC and NG contributed to the discussion of the results and to the proofreading of the final manuscript.

DP supervised EEG preprocessing and analysis, contributed to the discussion of the results and to the writing of the final manuscript.

